# Structure of the lysosomal SCARF (L-SCARF) complex, an Arf GAP haploinsufficient in ALS and FTD

**DOI:** 10.1101/2020.04.15.042515

**Authors:** Ming-Yuan Su, Roberto Zoncu, James H. Hurley

**Affiliations:** Department of Molecular and Cell Biology and California Institute for Quantitative Biosciences, University of California, Berkeley, Berkeley, CA 94720

## Abstract

Mutation of *C9ORF72* is the most prevalent defect in amyotrophic lateral sclerosis (ALS) and frontal temporal degeneration (FTD). Together with hexanucleotide repeat expansion, haploinsufficiency of *C9ORF72* contributes to neuronal dysfunction. We determined the structure of the SMCR8-C9orf72-WDR41 complex by cryo-EM. C9orf72 and SMCR8 are both longin-DENN domain proteins, while WDR41 is a beta-propeller protein that binds to SMCR8 such that the whole structure resembles an eye slip hook. Contacts between WDR41 and SMCR8^DENN^ drive lysosomal localization in amino acid starvation. The structure suggested that SMCR8-C9orf72 was a small GTPase activating protein (GAP). We found that SMCR8-C9orf72-WDR41 is a GAP for Arf family small GTPases, and refer to it as the Lysosomal SMCR8-C9orf72 Arf GAP (“L-SCARF”) complex. These data rationalize the function of C9orf72 both in normal physiology and in ALS/FTD.

## Main

Expansion of hexanucleotide GGGGCC repeats in the first intron of *C9ORF72* is the most prevalent genetic cause of amyotrophic lateral sclerosis (ALS) and frontal temporal degeneration (FTD) in humans^1,2^, accounting for approximately 40% of familial ALS, 5% of sporadic ALS and 10-50 % of FTD^3^. Two hypotheses, not mutually exclusive, have been put forward to explain how the mutation leads to progressive loss of neurons. The toxic gain of function hypothesis suggests that toxic molecules, including RNA and dipeptide repeat aggregates, disrupt neural function and lead to their destruction^4-14^. The loss of function hypothesis is based on the observation of a reduction in C9orf72 mRNA and protein levels in patients. In powerful support of the latter, the endogenous function of C9orf72 is essential for microglia ^15^ and for normal axonal actin dynamics in motor neurons ^16^, and restoring normal C9orf72 protein expression rescues function in *c9orf72* model neurons^17^.

C9orf72 is a longin and DENN (differentially expressed in normal and neoplastic cells) domain-containing protein^18^ (Fig. 1a). C9orf72 exists in cells as stable complex with another longin and DENN-containing protein, Smith-Magenis syndrome chromosome region, candidate 8 (SMCR8), and the WD repeat-containing protein 41 (WDR41)^19-24^ (Fig. 1a). For reasons described below, we will refer to SMCR8-C9orf72 as the “SCARF” complex. The main role of WDR41 appears to be to target SCARF to lysosomes^25^ via an interaction with the transporter PQ loop repeat-containing 2 (PQLC2)^26^. We therefore refer to SMCR8-C9orf72-WDR41 as the “L-SCARF” complex. Various cellular functions of SCARF in normal physiology have been proposed, including regulation of Rab-positive endosomes^27^, regulation of Rab8a and Rab39b in membrane transport^19,23^, regulation of the ULK1 complex in autophagy^20,23,24,28^, and regulation of mTORC1 at lysosomes^21,22,29^. Thus far it has been difficult to deconvolute which of these roles are direct *vs*. indirect. In order to gain more insight, we reconstituted and purified the complex, determined its structure, and assessed its function as a purified complex.

**Fig. 1:**
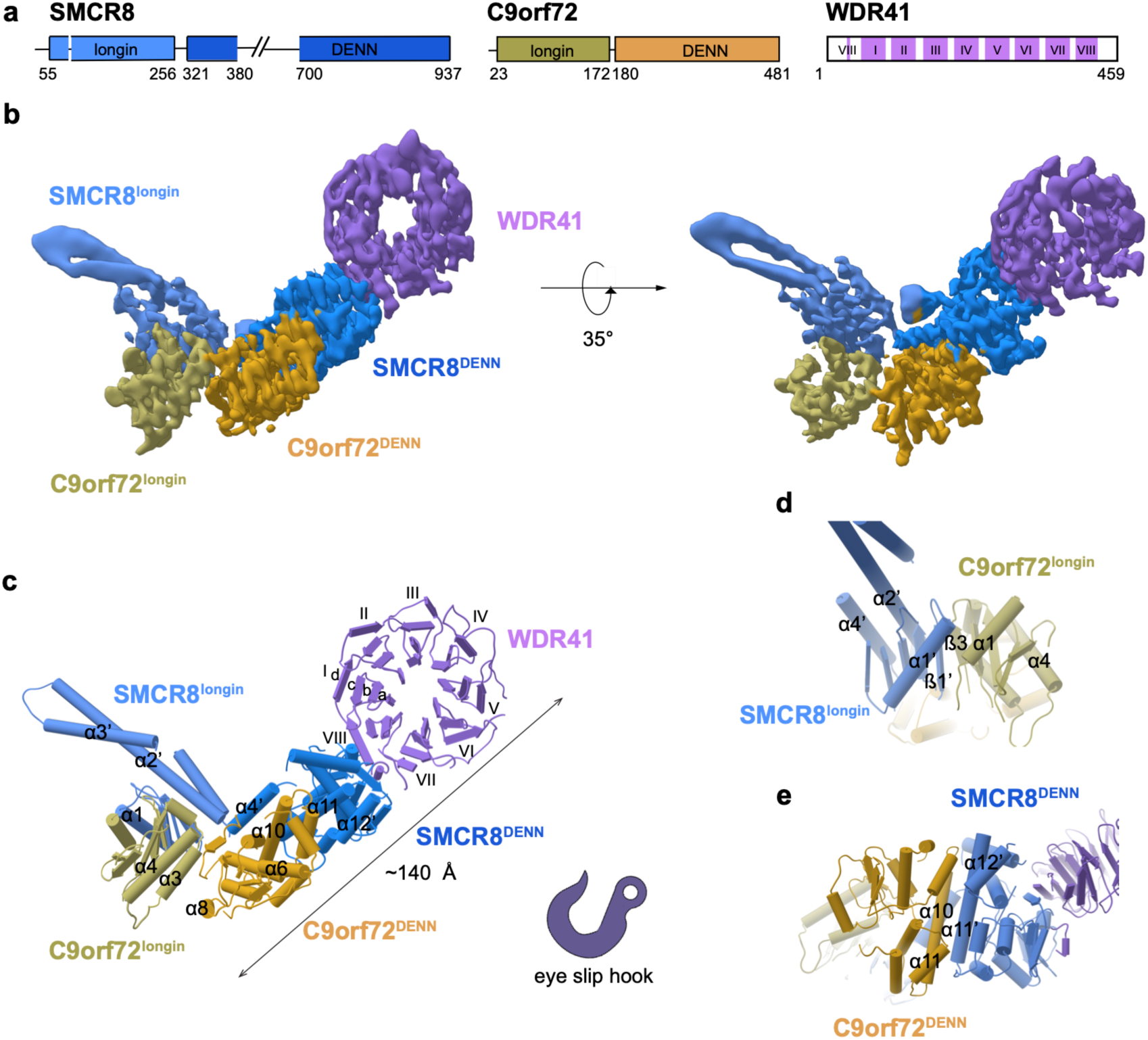
Cryo-EM structure of L-SCARF complex. **a**, Schematic diagram of the domain structure of L-SCARF complex. **b**, Cryo-EM density map (localfilter map, b-factor -50 Å^2^) and **c**, the refined coordinates of the complex shown as pipes and planks for α-helices and β-sheets, respectively. The domains color-coded as follows: SMCR8^longin^, cornflower blue; SMCR8^DENN^, dodger blue; C9orf72^longin^, olive; C9orf72^DENN^, goldenrod; WDR41, medium purple. Organizations of **d**, SMCR8^longin^: C9orf72^longin^ and **e**, SMCR8^DENN^: C9orf72^DENN^ arrangement.

Full length human SCARF and L-SCARF were generated in HEK293 Gn-Ti cells by simultaneous transient transfection of all subunits, and purified (Extended Data Fig. 1). The structure of L-SCARF was determined at a resolution of 3.8 Å by cryo-electron microscopy (cryo-EM) (Fig. 1b-c, Extended Data Fig. 2-4, Table 1). We were able to visualize the ordered ∼120 kDa portion of the complex, corresponding to about 60 % of the total mass of the complex. The structure has the shape of an eye slip hook with a long dimension of ∼140 Å (Fig. 1c). The ring of the hook was straightforward to assign to WDR41 by its appearance as an eight-bladed propeller. The remainder of the density evidenced two longin domains at the tip of the hook, with the bulk of the hook made up of two DENN domains. The SMCR8^DENN^ domain is in direct contact with WDR41, whilst C9orf72 has no direct contact with WDR41. The hook tip portion of the SMCR8^longin^ domain was assigned to residues S159-T210, which were predicted to comprise a long helical extension unique to SMCR8^longin^. SMCR8^longin^ and SMCR8^DENN^ are near each other but not in direct contact, and are connected by a helical linker consisting of residues T321-K363. Both domains of C9orf72 are positioned between SMCR8^longin^ and SMCR8^DENN^. This linear arrangement of domains gives the overall complex an elongated shape.

To map WDR41 interactions and facilitate interpretation of the cryo-EM structure, SCARF and L-SCARF complexes were subjected to hydrogen deuterium exchange mass spectrometry (HDX-MS) for 0.5, 5, 50, 500 and 50,000 sec and compared to each other (Fig. 2, Extended Data Fig. 1, 5, 6, Dataset 1). Excellent peptide coverage (89, 87 and 80 % for SMCR8, C9orf72 and WDR41, respectively) was achieved and consistent patterns were observed at all experimental time points. Several regions in SMCR8 including the N-terminal 54 residues, residues V104-V118, E212-I230, P257-F315, V378-I714 and V788-Y806 showed more than 50 % deuterium uptake at 0.5 sec, indicating these regions are intrinsically disordered regions (IDRs), consistent with sequence-based predictions. Nearly all of C9orf72 was protected from exchange, except for the N-terminal 21 residues and the C-terminus. For WDR41, the N-terminal 24 residues, and the loops connecting blade II-III (R128-C131), blade V-VI (R260-D270, L277-I284), internal loop of blade VII and the loop connecting to VIII (R352-L357, M369-E396) were flexible.

**Fig. 2:**
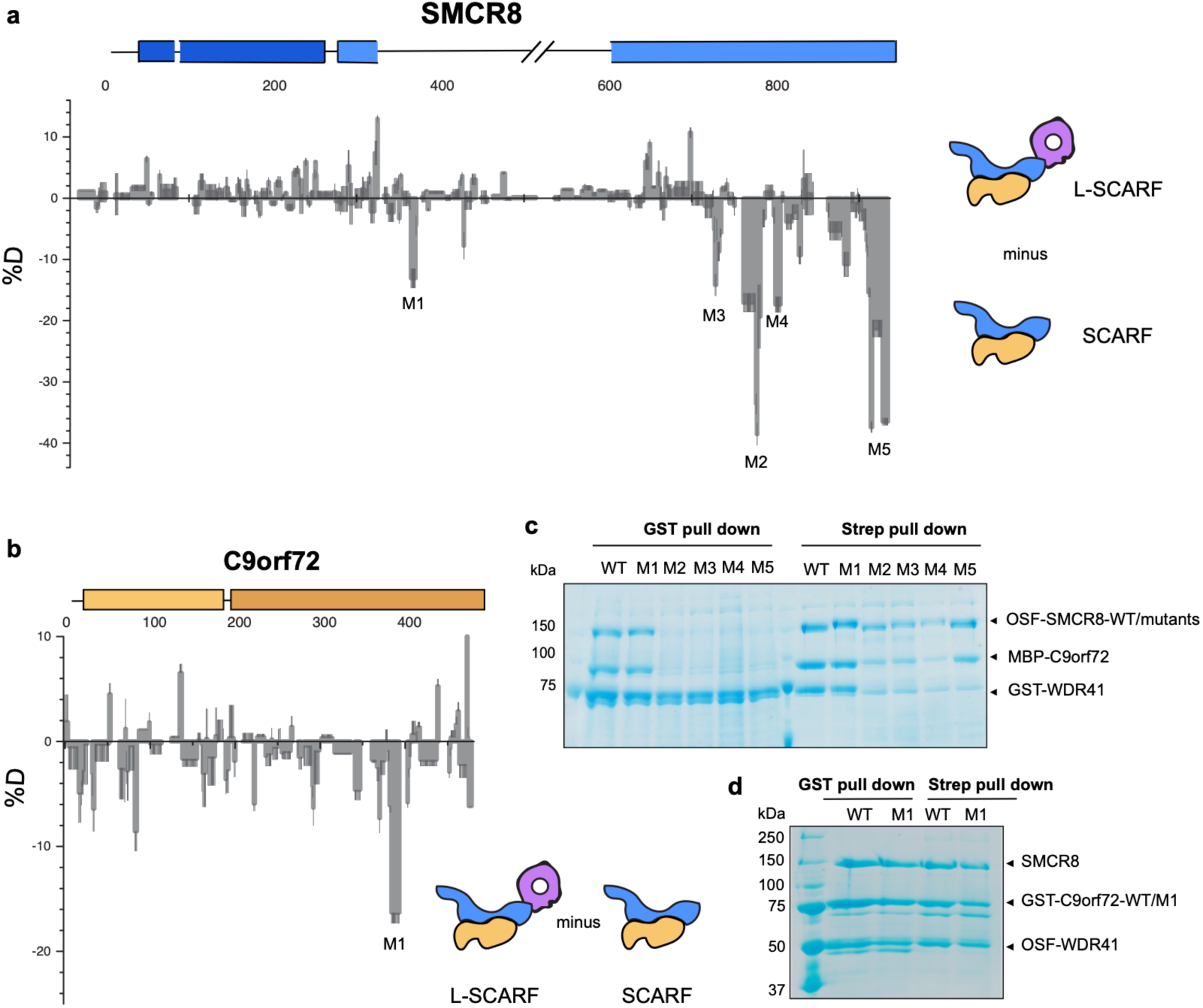
HDX-MS of SCARF in the absence of WDR41. **a**, Difference plot of percentage of deuteron incorporation of SMCR8 in heterotrimer versus dimer at 5 sec timepoint. **b**, Difference plot of percentage of deuteron incorporation of C9orf72 in heterotrimer versus dimer at 0.5 sec timepoint. **c**, Pull down experiment of SMCR8 mutants with wild type C9orf72 and WDR41. **d**, Pull down experiments of C9orf72 mutant with wild type SMCR8 and WDR41.

Difference heat maps for C9orf72 and SMCR8 (Fig. 2a-b) showed that in presence of WDR41, regions of the SMCR8^DENN^ including K363-L372 (SMCR8^M1^), P763-Q770 (SMCR8^M2^), S729-V735 (SMCR8^M3^), T807-D811 (SMCR8^M4^) and C-terminal K910-Y935 (SMCR8^M5^) were protected from exchange (Fig. 2, Extended Data Fig. 5, 6), consistent with the structure. There was no significant change in C9orf72, with the exception of K388-R394 (C9orf72^M1^) (Fig. 2). Regions showing protection changes were mutagenized and tested in pull down experiments (Fig. 2c-d). Except for the helical linker mutant SMCR8^M1^, the mutations including SMCR8^M2-M5^ abolished the interaction with WDR41. When WDR41 failed to pull down SMCR8 mutants, wild-type C9orf72 was not detected either. This confirms the structural finding that SMCR8 bridges the other two components. Because C9orf72^M1^ retained interaction with SMCR8-WDR41, we concluded that this region was protected by a conformational change induced upon WDR41 binding, consistent with the lack of direct interaction in the cryo-EM structure. The cryo-EM structure showed that SMCR8 bound to blades VIII and C terminal helix of WDR41 (Fig. 3a-b). The pull down experiment showed that the N-terminal residues E35-K40 of blade VIII and the C-terminal helix S442-V459 are required for SMCR8 binding (Extended Data Fig.7). Collectively, the HDX-MS and mutational results corroborate the structural interpretation.

**Fig. 3:**
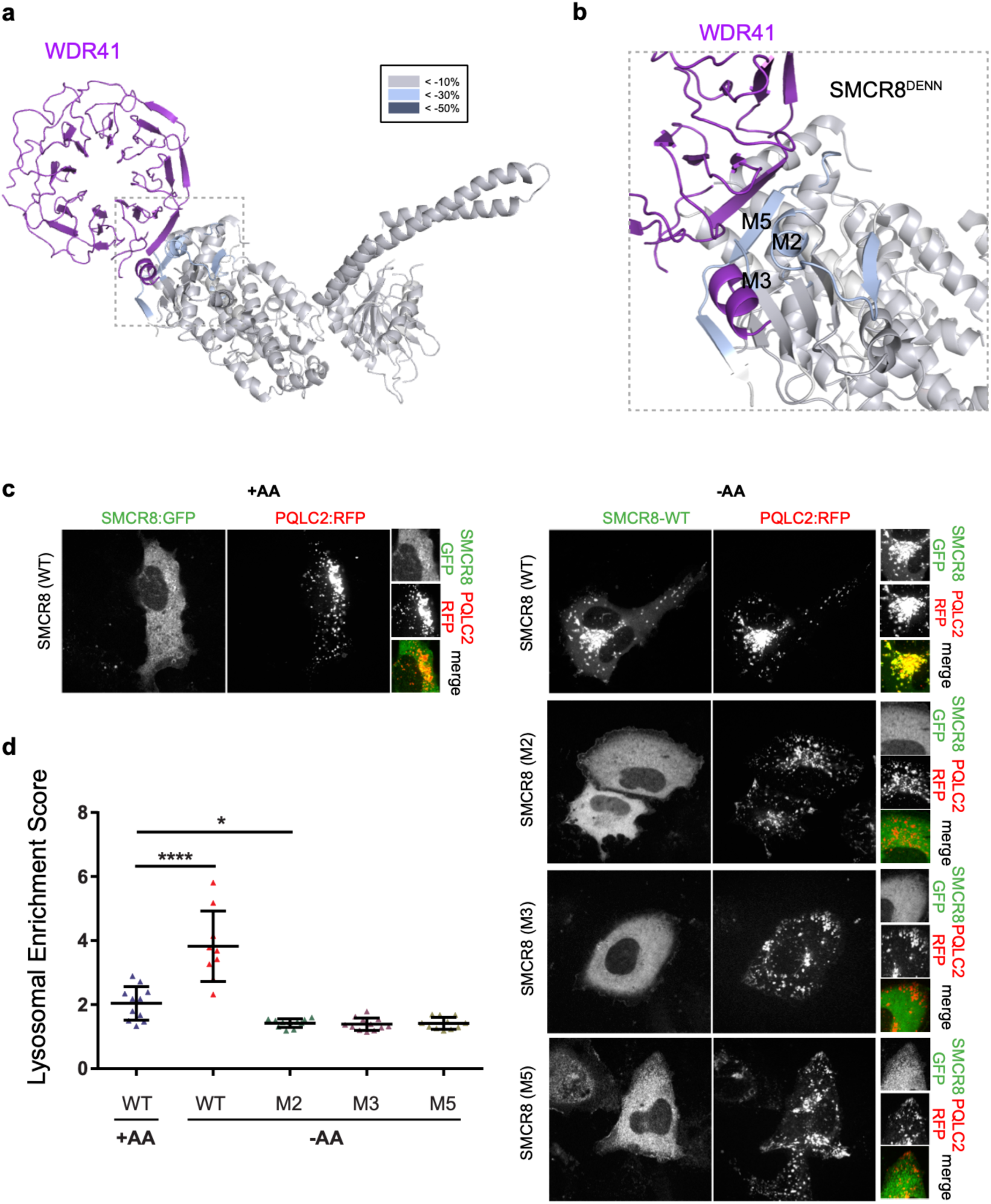
SMCR8 mutants fail to localize on lysosome. **a**, HDX uptake difference at 0.5 sec was mapped on SCARF. **b**, Close view of SMCR8-WDR41 interface, highlighting the SMCR8 mutants. **c**, SMCR8-PQLC2 lysosome colocalization experiment in cells expressing the indicated SMCR8 constructs under the indicated nutrient conditions. **d**, Quantification of SMCR8 lysosomal enrichment score for immunofluorescence images in **c**. More than 10 cells were quantified for each condition.

WDR41 is responsible for the reversible targeting of SCARF to lysosomes in nutrient depletion^25^. WDR41 in turns binds to lysosomes via PQLC2^26^. We co-transfected DNA encoding GFP-SMCR8, C9orf72, WDR41 and PQLC2-mRFP in HEK293A cells. SMCR8 clustered on PQLC2-positive lysosomes in amino acid depletion and was diffusely localized in the cytosol upon refeeding (Fig. 3c), consistent with these reports^25,26^. SMCR8 mutants deficient in WDR41 binding *in vitro* did not colocalize with PQLC2-postive lysosomes, but rather were diffusely localized in the cytosol even under amino acid-starved conditions (Fig. 3c-d). These findings confirm that the WDR41 binding site on SMCR8 as mapped by cryo-EM and HDX-MS is responsible for the lysosomal localization of the complex in amino-acid starvation.

The structure showed that SMCR8^longin^ forms a heterodimer with C9orf72^longin^ in the same manner as Nprl2-Nprl3 of the GATOR1 complex^30^ and FLCN-FNIP2 in the Lysosomal Folliculin Complex (LFC)^31,32^. The Nprl2 and FLCN subunits of these complexes are the GTPase activating proteins (GAPs) for the lysosomal small GTPases RagA^33^ and RagC^34^, respectively. Structure-based alignment of SMCR8 with FLCN and Nprl2 showed they shared a conserved Arg finger residue^31,32,35^ (Fig. 4a), corresponding to SMCR8 Arg147. This Arg residue is exposed on the protein surface near the center of a large concave surface that appears suitable for binding a small GTPase (Extended Data Fig.8). Using a Trp fluorescence-based assay, we assayed SCARF for GAP activity with respect to RagA or RagC and found none detectable (Extended Data Fig. 9). We also assayed for GAP activity with respect to Rab1a^28^ and the late endosomal Rab7^27^, and again, activity was undetectable (Extended Data Fig. 9,10).

**Fig. 4:**
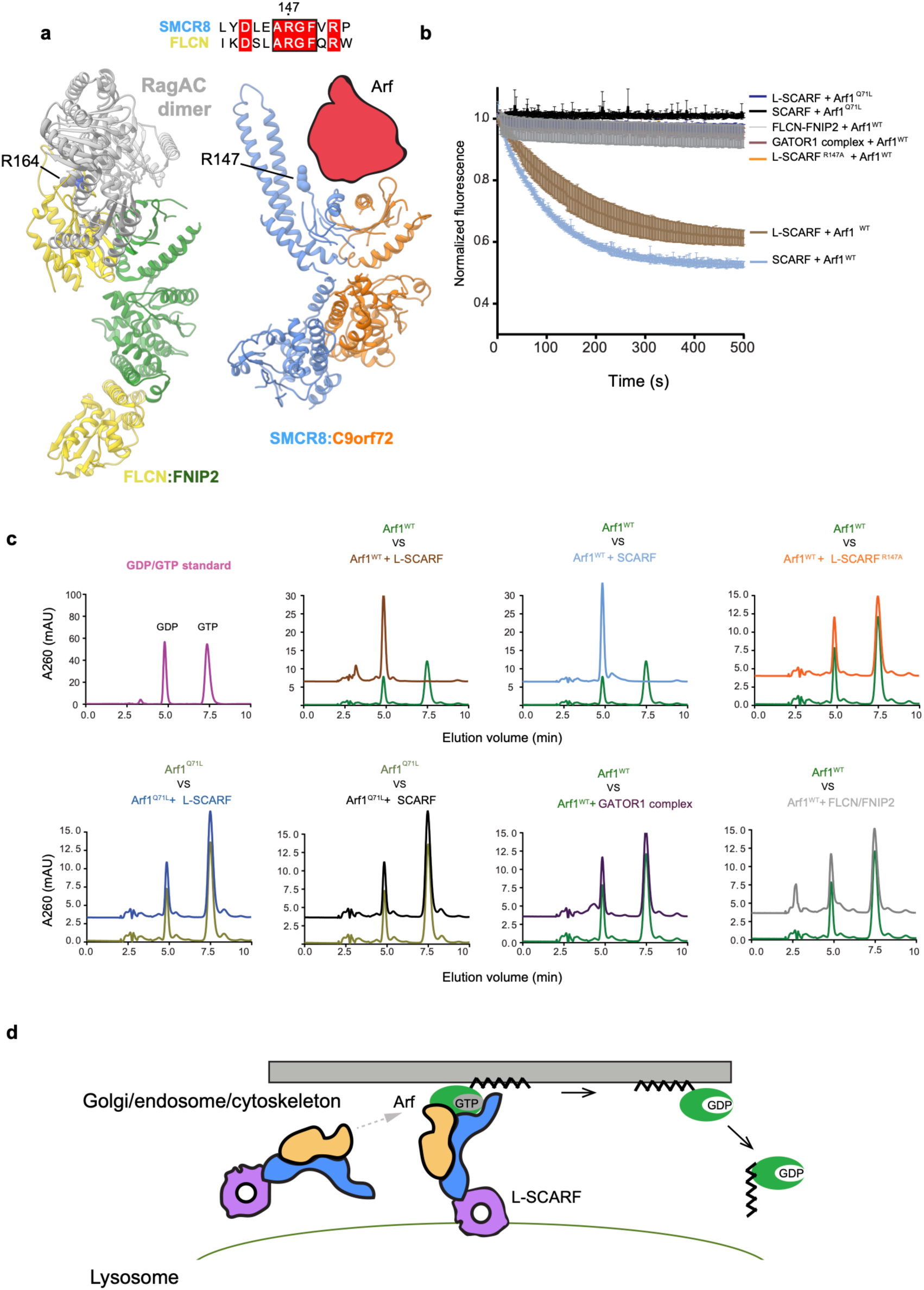
SCARF is a GAP for Arf proteins. **a**, Structure comparison of FNIP2-FLCN and SCARF, implying a potential binding site for substrates. The conserved Arg residue was shown in spherical representation. **b**, Tryptophan fluorescence GTPase signal was measured for Arf1^WT or Q71L^ before and after addition of SCARF^WT or R147A^ -WDR41, SCARF, FLCN-FNIP2 or GATOR1 complex. **c**, HPLC-based GTPase assay with Arf1^WT or Q71L^ proteins in the absence and addition of GAP complex as indicated. **d**, Model for Arf protein family activation by SCARF-WDR41.

It has been reported that C9orf72 interacts with the small GTPases Arf1 and Arf6^36^ in neurons^16^, although the nature of the interaction is unknown. We found that L-SCARF was an efficient GAP for Arf1 on the basis of both Trp fluorescence and HPLC-based assays (Fig. 4). The Arf1^Q71L^ GTP locked mutant had no activity (Fig.4b-c), nor did the version of the complex containing the SMCR8^R147A^ finger mutation. FLCN-FNIP2 and GATOR1 had no GAP activity towards Arf1. SCARF was as active as L-SCARF, consistent with the location of WDR41 on the opposite side of the complex from Arg147. L-SCARF has activity against the other Arf family members, Arf5 and Arf6, but not against the lysosomal Arf-like proteins Arl8a and Arl8b (Extended Data Fig. 9, 10). These observations clarify the nature of the reported C9orf72-Arf interaction by showing that the role of C9orf72 is to stabilize a complex with SMCR8, which is in turn an efficient and selective GAP for Arf GTPases. For this reason, we have adopted the term SCARF for <u>S</u>MCR8-<u>C</u>9orf72 <u>ARF</u> GAP for the complex, and L-SCARF for the WDR41-containing version that is lysosomally localized in amino acid starvation.

These structural and functional data shed light on the normal function of C9orf72, which is thought to contribute to neuronal loss of function in ALS and FTD^17^. The structure shows that C9orf72 is the central component of its complex with SMCR8. The longin and DENN domains of SMCR8 flank and are stabilized by C9orf72. SMCR8 contains the binding site for WDR41 that is responsible for lysosomal localization during amino acid starvation. The structure shows that SCARF belongs to the same class of double-longin domain GAP complexes as GATOR1^30^ and FLCN-FNIP^31,32^. Unlike GATOR1 and FLCN-FNIP, SCARF is inactive against Rag GTPases, but is active against Arf GTPases instead. The GAP active site is located at the opposite end of the complex from the lysosomal targeting site on WDR41.

A remaining question concerns the regulation of the Arf-GAP function of SMCR8-C9orf72 in cells. Our *in vitro* observation that SCARF and L-SCARF have comparable GAP activities suggests that, in cells, SMCR8-C9orf72 may regulate Arf GTPases both in full nutrient conditions, when the complex is primarily localized in the cytosol, and under amino acid starvation, when it relocalizes to the lysosomal membrane via WDR41-PQLC2 interaction. However, additional factors could limit or augment the Arf-GAP activity in either condition and restrict or enhance access to the GTP-bound Arf substrate. Arf proteins are not observed on lysosomes, and their closest lysosomal cousins, Arl8a and Arl8b, are not substrates for SCARF. Thus, sequestration of L-SCARF on lysosomes could prevent it from regulating the Arfs in *cis* under unfavorable metabolic conditions. Alternatively, L-SCARF could act in *trans* on Arf bound to the membrane of a compartment other than the lysosome. Arf GTPases are found on the Golgi, endosomes, plasma membrane, cytoskeleton, and in the cytosol^36^, and typically function on membranes in their active GTP-bound form. Several reports have found C9orf72 to be associated with endosomes^17,27,37^ and the cytoskeleton^16^, which are good candidates for the locus of the Arf substrate of SCARF. The potential *trans* GAP activity of L-SCARF *vs*. endosomal or cytoskeletal Arf would be facilitated by its elongated structure and the distal positioning of the GAP and lysosomal localization sites (Fig. 4d).

The structure of the complex places it in the same class of heterodimeric longin-DENN domain protein GAP complexes as FLCN-FNIP^31,32^. FLCN-FNIP is a major node for communicating lysosomal nutrient status to the nucleus by virtue of its regulation of the RagC GTPase^34^ and the phosphorylation of transcription factors regulating lysosome biogenesis and autophagy^31^. These data show how C9orf72 is stabilized on the lysosome in amino acid starvation by bridging contacts made by SMCR8 to WDR41. They show that C9orf72 serves as the central hub that stabilizes the SCARF complex, in which SMCR8 is the “business end” with respect to Arf GAP activity. A number of physiological functions have been imputed to C9orf72 based on protein interactions mapped in cells and in lysates, but it has been unclear what biochemical activities belong to C9orf72 itself, as opposed to downstream and indirect effects. Here, we established a direct function for purified SMCR8-C9orf72, which we designate the SCARF complex for its robust and specific GAP activity on Arf proteins. We found that this activity is comparable to that of other well-established GAP complexes such as GATOR1 and FLCN-FNIP with respect to their substrates. This activity likely explains how C9orf72 modulates actin dynamics in neurons^16^. It has been reported that Arf1 promotes mTORC1 activation^38^, so the Arf GAP function of SCARF could explain how this complex antagonizes mTORC1^29^. Finally, multiple reports connect C9orf72 to endosomal sorting^17,27,37^, a process in which the role of Arfs is well-established^36^. The structural and *in vitro* biochemical data reported here thus provide a framework and a foothold for understanding how the normal functions of C9orf72 relate to lysosomal signaling, autophagy, and neuronal survival.

## Supporting information

Extended Data Set 1 HDX-MS

## Methods

### Protein expression and purification

Synthetic genes encoding SMCR8 were amplified by PCR and cloned into the pCAG vector coding for an N-terminal twin-STREP-FLAG tag using KpnI and XhoI restriction sites. The pCAG vector encoding an N-terminal GST followed by a TEV restriction site or uncleaved MBP tag was used for expression of C9orf72. WDR41 was cloned into pCAG vector without a tag or with a GST tag for pull down experiments. For the mutations of SMCR8 identified from HDX experiments, SMCR8^M1^ (K363-L371) was mutated to MSDYDIPTTE, which is a 10-residue linker derived from the pETM11 vector. SMCR8^M2^ (P771-Q778) or (K762-L782) for lysosome localization experiments was mutated to GGKGSGGS. SMCR8^M3^ (S729-V735) and SMCR8^M4^ (T807-D811) were made by mutating these regions to GGKGSGG and GGKGS, respectively. SMCR8^M5^ was made by truncation after residue 910K. C9orf72^M1^ (K388-L393) was mutated to polyAla. The SMCR8 arginine finger mutation R147A was made using two step PCR and cloned into the expression vector.

HEK293-GnTi cells adapted for suspension were grown in Freestyle media supplemented with 1% FBS and 1% antibiotic-antimycotic at 37 °C, 80 % humidity, 5 % CO2, and shaking at 140 rpm. Once the cultures reached 1.5–2 million cells mL−1 in the desired volume, they were transfected as followed. For a 1 L transfection, 3 mL PEI (1 mg ml−1, pH 7.4, Polysciences) was added to 50 mL hybridoma media (Invitrogen) and 1 mg of total DNA (isolated from transformed E. coli XL10-gold) in another 50 mL hybridoma media. 1 mg of transfection DNA contained equal mass ratio of C9orf72 complex expression plasmids. PEI was added to the DNA, mixed and incubated for 15 min at room temperature. 100 mL of the transfection mix was then added to each 1 L culture. Cells were harvested after 3 days.

Cells were lysed by gentle rocking in lysis buffer containing 50 mM HEPES, pH 7.4, 200 mM NaCl, 2 mM MgCl_2_, 1% (vol/vol) Triton X-100, 0.5 mM TCEP, protease inhibitors (AEBSF, Leupeptin and Benzamidine) and supplemented with phosphatase inhibitors (50 mM NaF and 10 mM beta-glycerophosphate) at 4 °C. Lysates were clarified by centrifugation (15,000 g for 40 min at 4 °C) and incubated with 5 mL glutathione Sepharose 4B (GE Healthcare) for 1.5 hr at 4 °C with gentle shaking. The glutathione Sepharose 4B matrix was applied to a gravity column, washed with 100 mL wash buffer (20 mM HEPES, pH 7.4, 200 mM NaCl, 2 mM MgCl_2_, and 0.5 mM TCEP), and purified complexes were eluted with 40 mL wash buffer containing 50 mM reduced glutathione. Eluted complexes were treated with TEV protease at 4 °C overnight. TEV-treated complexes were purified to homogeneity by injection on Superose 6 10/300 (GE Healthcare) column that was pre-equilibrated in gel filtration buffer (20 mM HEPES, pH 7.4, 200 mM NaCl, 2 mM MgCl_2_, and 0.5 mM TCEP). For long-term storage, fractions from the gel filtration chromatography were frozen using liquid nitrogen and kept at -80 °C. SCARF and L-SCARF were expressed and purified using the same protocol.

For expression of His_6_-tagged Arf1 (residue E17-K181), Arf1 Q71L, Arf5 (residue Q17-Q180), Arf6 (residue R15-S175), Arf6 Q67L, His_6_-Rab1a, His_6_-Arl8a (E20-S186) and His_6_-Arl8b (E20-S186) proteins, plasmids were transformed into *E*.*coli* BL21 DE3 star cells and induced with 0.5 mM IPTG at 18° C overnight. The cells were lysed in 50 mM Tris-HCl pH 8.0, 300 mM NaCl, 2 mM MgCl_2_, 5 mM imidazole, 0.5 mM TCEP and 1 mM PMSF by ultrasonication. The lysate was centrifuged at 15,000 g for 30 min. The supernatant was loaded into Ni-NTA resin and washed with 20 mM imidazole and eluted with 300 mM imidazole. The eluate was further purified on a Superdex 75 10/300 (GE Healthcare) column equilibrated in 20 mM HEPES, pH 7.4, 200 mM NaCl, 2 mM MgCl_2_, and 0.5 mM TCEP. Rag, FLCN-FNIP2 and GATOR1 complex were purified as described previously^31^. GST-tagged Rab7 was expressed in the same conditions as above and purified with GST resin, eluted in 50 mM reduced glutathione and applied on Superdex 200 column.

### Hydrogen/Deuterium exchange experiment

Sample quality was assessed by SDS-PAGE before each experiment. Amide hydrogen exchange mass spectrometry was initiated by a 20-fold dilution of 10 µM L-SCARF or SCARF into 95 µl D_2_O buffer containing 20 mM HEPES pH (pD 8.0), 200 mM NaCl, 1 mM TCEP at 30° C. Incubations in deuterated buffer were performed at intervals from 0.5, 5, 50, 500 and 50,000 sec (0.5 sec was carried out by incubating proteins with ice cold D_2_O for 5 sec). All exchange reactions were carried out in triplicate or quadruplicate. Backbone amide exchange was quenched at 0° C by the addition of ice-cold quench buffer (400 mM KH_2_PO_4_/H_3_PO_4_, pH 2.2). The 50,000 sec sample served as the maximally labeled control. Quenched samples were injected onto a chilled HPLC setup with in-line peptic digestion and then eluted onto a BioBasic 5 µM KAPPA Capillary HPLC column (Thermo Fisher Scientific), equilibrated in buffer A (0.05 % TFA), using 10-90 % gradient of buffer B (0.05 % TFA, 90 % acetonitrile) over 30 mins. Desalted peptides were eluted and directly analyzed by an Orbitrap Discovery mass spectrometer (Thermo Fisher Scientific). The spray voltage was 3.4 kV and the capillary voltage was 37 V. The HPLC system was extensively cleaned between samples. Initial peptide identification was performed via tandem MS/MS experiments. A Proteome Discoverer 2.1 (Thermo Fisher Scientific) search was used for peptide identification and coverage analysis against entire complex components, with precursor mass tolerance ± 10 ppm and fragment mass tolerance of ± 0.6 Da. Mass analysis of the peptide centroids was performed using HDExaminer (Sierra Analytics), followed by manual verification of each peptide.

### Cryo-EM grid preparation and data acquisition

The purified L-SCARF complex was diluted to 0.8 µM in 20 mM HEPES pH 7.4, 2 mM MgCl_2_, and 0.5 mM TCEP and applied to glow-discharged C-flat (1.2/1.3, Au 300 mesh) grids. The sample was vitrified after blotting for 2 sec using a Vitrobot Mark IV (FEI) with 42 sec incubation, blot force 8 and 100 % humidity. The complex was visualized with a Titan Krios electron microscope (FEI) operating at 300 kV with a Gatan Quantum energy filter (operated at 20 eV slit width) using a K2 summit direct electron detector (Gatan, Inc.) in super-resolution counting mode, corresponding to a pixel size of 0.5745 Å on the specimen level. In total, 3,508 movies were collected in nanoprobe mode using Volta phase plate (VPP) with defocus collected around -60 nm. Movies consisted of 49 frames, with a total does of 59.8 e-/Å^2^, a total exposure time of 9.8 sec, and a dose rate of 8.1 e^-^/pixel/sec. Data were acquired with SerialEM using custom macros for automated single particle data acquisition. Imaging parameters for the data set are summarized in Extended Data Table 1.

### Cryo-EM data processing

Preprocessing was performed during data collection within Focus^39^. Drift, beam induced motion and dose weighting were corrected with MotionCor2^40^ using 5 × 5 patches. CTF fitting and phase shift estimation were performed using Gctf v1.06 ^41^, which yielded the characterized pattern of phase shift accumulation over time for each position. The data were manually inspected and micrographs with excess ice-contamination or shooting on the carbon were removed. A total of 4,810,184 particles from 3,220 micrographs were picked using gautomatch (http://www.mrc-lmb.cam.ac.uk/kzhang/Gautomatch/) and extracted with binning 4. All subsequent classification and reconstruction steps were performed using Relion3-beta^42^ or cryoSPARC v2 ^43^. The particles were subjected to 3D classification (K=5) using a 60 Å low-pass filtered *ab initio* reference generated in cryoSPARC. Around 2.2 million particles from the two best classes were selected for 3D auto-refinement and another round of 3D classification (K=8, T=8, E-step=8) without alignment. Some 1.8 millions particles from the best 6 classes were reextracted with binning 2 and refined to 4.9 Å, and further subjected to 2D classification without alignment for removing contamination and junk particles. After another round of 3D classification (K=4) with alignment, the best class was extracted and imported into cryoSPARC v2 for another round of 2D classification. The cleaned up 571,002 particles were applied to CTF refinement, Bayesian polishing and further particles at edges were removed in Relion 3. Final 381, 450 particles resulted in final resolution of 3.8 Å with a measured map B-factor of -102 Å^2^. More extensive 3D classification, focus classification in Relion3 did not improve the quality of the reconstruction. Local filtering and B-factor sharpening were done in cryoSPARC v2. All reported resolutions are based on the gold-standard FSC 0.143 criterion.

### Atomic model building and refinement

The model of WDR41 was generated with I-Tasser^44^ and used 5nnz, 2ymu, 5wlc, 4nsx and 6g6m as starting models. The model of the C9orf72^longin^ domain was generated based on the Nprl2^longin^ domain (pdb 6ces) in Modeller^45^. The model of SMCR8^DENN^ domain was generated from Modeller and RaptorX^46^ using the FLCN^DENN^ domain (pdb 3v42) or the *DENN*<u>D1B</u> ^DENN^ domain (pdb 3tw8) as templates. The SMCR8^longin^ and C9orf72^DENN^ domain were generated with Phyre2^47^ using FLCN^longin^ and FNIP2^DENN^ domain (pdb 6nzd) as templates. Secondary structure predictions of each protein were carried out with Phyre2^47^ or Psipred^48^. The models were docked into the 3D map as rigid bodies in UCSF Chimera ^49^. The coordinates of the structures were manually adjusted and rebuilt in Coot^50^. The resulting models were refined using Phenix.real_space.refine in the Phenix suite with secondary structure restraints and a weight of 0.1^51,52^. Model quality was assessed using MolProbity^53^ and the map-*vs*-model FSC (Extended Data Table 1 and Extended Data Fig. 4a). Data used in the refinement excluded spatial frequencies beyond 4.2 Å to avoid over fitting. A half-map cross-validation test showed no indication of overfitting (Extended Data Fig. 4b). Figures were prepared using UCSF Chimera ^49^ and PyMOL v1.7.2.1. The cryo-EM density map has been deposited in the Electron Microscopy Data Bank under accession code EMD-21048 and the coordinates have been deposited in the Protein Data Bank under accession number 6V4U.

### Live cell imaging

800,000 HEK 293A cells were plated onto fibronectin-coated glass-bottom Mattek dishes and transfected with the indicated wild type GFP-SMCR8 or mutants, C9orf72, WDR41 and PQLC2-mRFP constructs with transfection reagent Xtremegene. 24 hrs later, cells were starved for amino acids for one hr (-AA) or starved and restimulated with amino acids for 10 mins (+AA). Cells in the -AA condition were transferred to imaging buffer (10 mM HEPES, pH7.4, 136 mM NaCl, 2.5 mM KCl, 2 mM CaCl_2_, 1.2 mM MgCl_2_) and cells in the +AA condition were transferred to imaging buffer supplemented with amino acids, 5 mM glucose, and 1% dialyzed FBS (+AA) and imaged by spinning-disk confocal microscopy. Lysosomal enrichment was scored as described^31^ using a home-built Matlab script to determine the lysosomal enrichment of GFP SMCR8. The score was analyzed for at least ten cells for each condition. Unpaired t-tests were calculated using Prism 6 (Graphpad).

### HPLC analysis of nucleotides

The nucleotides bound to small GTPases were assessed by heating the protein to 95 °C for 5 min followed by 5 min centrifugation at 16,000 g. The supernatant was loaded onto a HPLC column (Eclipse XDB-C18, Agilent). Nucleotides were eluted with HPLC buffer (10 mM tetra-n-butylammonium bromide, 100 mM potassium phosphate pH 6.5, 7.5 % acetonitrile). The identity of the nucleotides was compared to GDP and GTP standards.

### HPLC-based GAP assay

HPLC-based GTPase assays were carried out by incubating 30 µl of GTPases (30 µM) with or without GAP complex at a 1:50 molar ratio for 30 min at 37 °C. Samples were boiled for 5 min at 95 °C and centrifuged for 5 min at 16,000 g The supernatant was injected onto an HPLC column as described above. The experiments are carried out in triplicate and one representative plot is shown.

### Tryptophan fluorescence-based GAP assay

Fluorimetry experiments were performed using a FluoroMax-4 (Horiba) instrument and a quartz cuvette compatible with magnetic stirring, a pathlength of 10 mm, and were carried out in triplicate. The Trp fluorescence signal was collected using 297 nm excitation (1.5 nm slit) and 340 nm emission (20 nm slit). Experiments were performed in gel filtration buffer at room temperature with stirring. Data collection commenced with an acquisition interval of 1 sec. 2 µM GTPase was added to the cuvette initially. Once the signal was equilibrated, SCARF^WT or R147A^-WDR41 or SCARF, FLCN-FNIP2, or GATOR1 complex was pipetted into the cuevette at a 1: 10 molar ratio. Time t = 0 corresponds to GAP addition. The fluorescence signal upon GAP addition was normalized to 1 for each experiment. Mean and standard error of the mean of three replicates per conditions were plotted.

## Acknowledgments

We thank D. Toso, S. Fromm, K. L. Morris, V. Kasinath and P. Tobias for cryo-EM advice and support, X. Shi for HDX support, S. Fromm for comments on the manuscript, C. Behrends and G. Stjepanovic for suggestions and contributions to the early stages of the project, and R. Lawrence for cell imaging advice. Access to the FEI Titan Krios was provided through the BACEM UCB facility. This work was supported by NIH grants R01GM111730 (J.H.H.) and R01GM130995 (R.Z.), the Bakar Fellows program (J.H.H.), the Pew-Stewart Scholarship for Cancer Research and Damon Runyon-Rachleff Innovation Award (R.Z.), and a postdoctoral fellowship from the Association for Frontotemporal Degeneration (M.-Y.S.).

## Author contributions

Conceptualization, M.-Y.S. and J.H.H.; Investigation, M.-Y.S. and R. Z.; Supervision, J.H.H., R.Z.; Writing-original draft, M.-Y.S. and J.H.H.; Writing-review and editing, all authors.

## Data availability

EM density map has been deposited in the EMDB with accession number EMD-21048. Atomic coordinates for the L-SCARF have been deposited in the PDB with accession number 6V4U.

## Extended data figures/table

**Extended Data Fig. 1:**
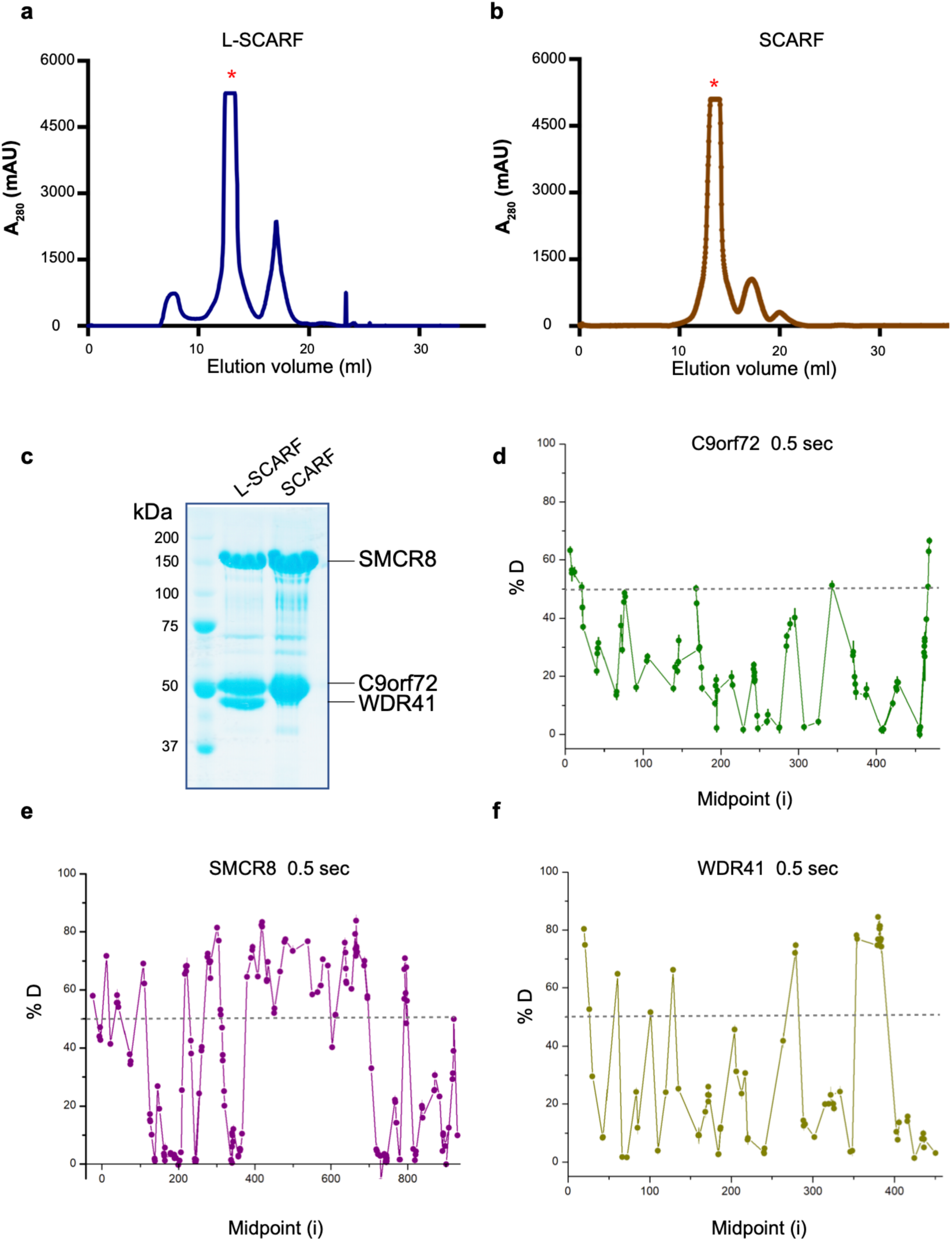
Purification of the L-SCARF and SCARF complex as well as the HDX data for trimer. **a**, The superose 6 gel filtration elution profile for L-SCARF complex. **b**, The superose 6 gel filtration elution profile for SCARF complex. mAU, milli-absorbance units. **c**, The purified full length L-SCARF and SCARF were analyzed by SDS-PAGE. **d-f**, Deuterium uptake data for L-SCARF complex at 0.5 sec timepoint with error bars from triplicate measurements. Peptides with more than 50 % deuterium uptake are the flexible regions. Y axis represents the average percent deuteration. X axis demonstrates the midpoint of a single peptic peptide.

**Extended Data Fig. 2:**
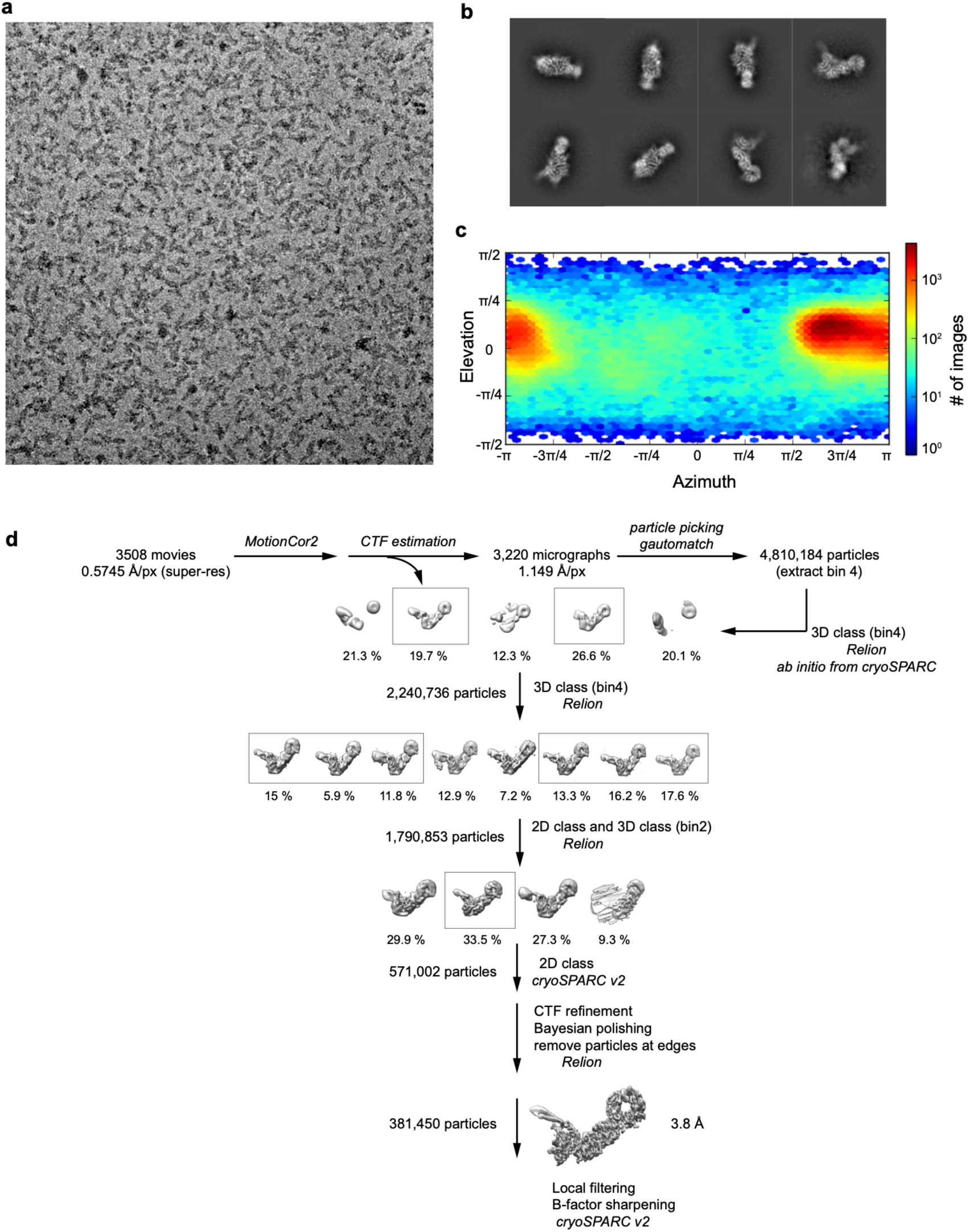
Cryo-EM data processing. **a**, Representative cryo-EM micrograph of L-SCARF complex. **b**, Representative 2D classes. **c**, Orientation distribution of the aligned particles. **d**, Image processing procedure.

**Extended Data Fig. 3:**
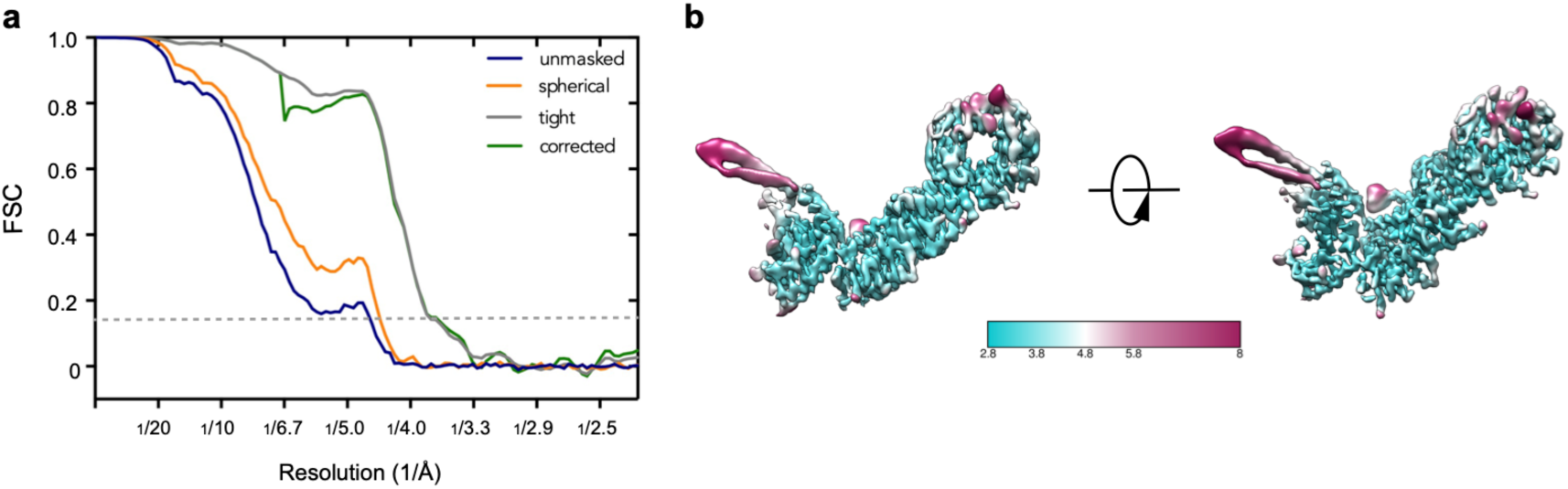
Resolution estimation of the cryo-EM map. **a**, Comparison between FSC curves. **b**, L-SCARF complex map color-coded by the local resolution estimation.

**Extended Data Fig. 4:**
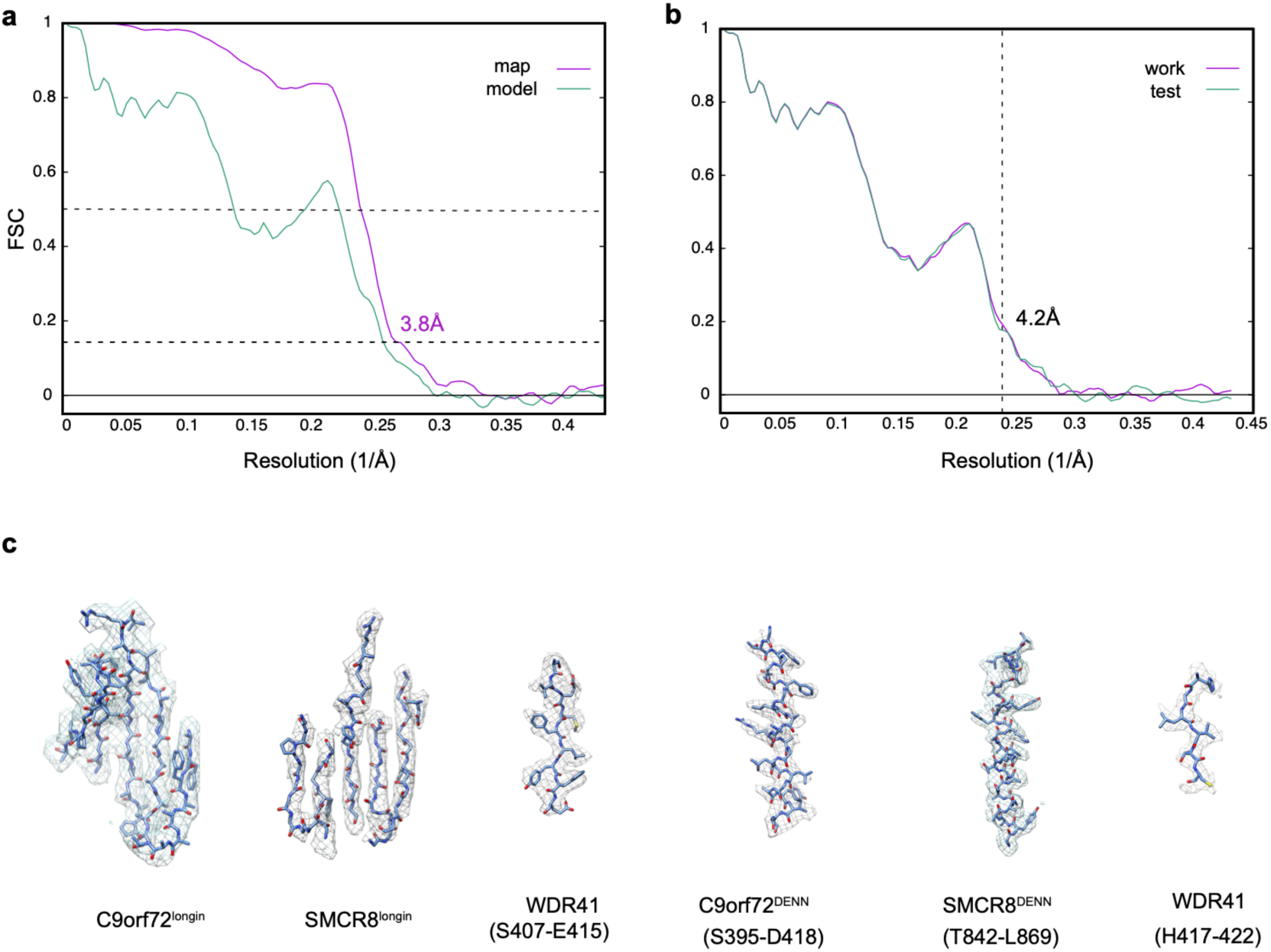
Model building and validation. **a**, Refinement and map-vs-model FSC. **b**, Cross-validation test FSC curves to assess overfitting. The refinement target resolution (4.2 Å) is indicated. **c**, Refined coordinate model fit of the indicated region in the cryo-EM density.

**Extended Data Fig. 5:**
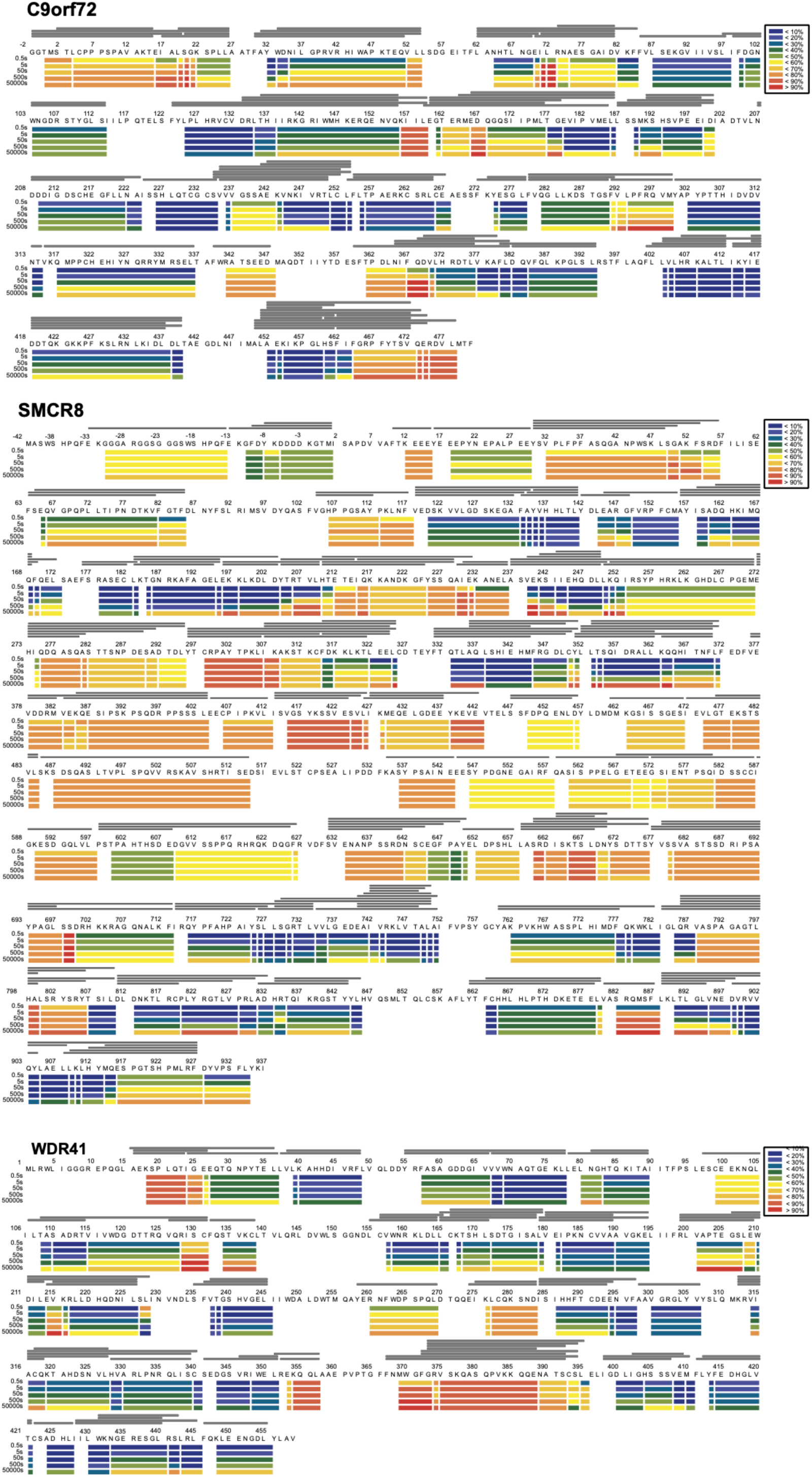
Deuterium uptake of L-SCARF complex. HDX-MS data are shown in heatmap format where peptides were represented using rectangular strips above the protein sequence. Absolute deuterium uptake after 0.5, 5, 50, 500 and 50,000 sec were indicated by a color gradient below the protein sequence.

**Extended Data Fig. 6:**
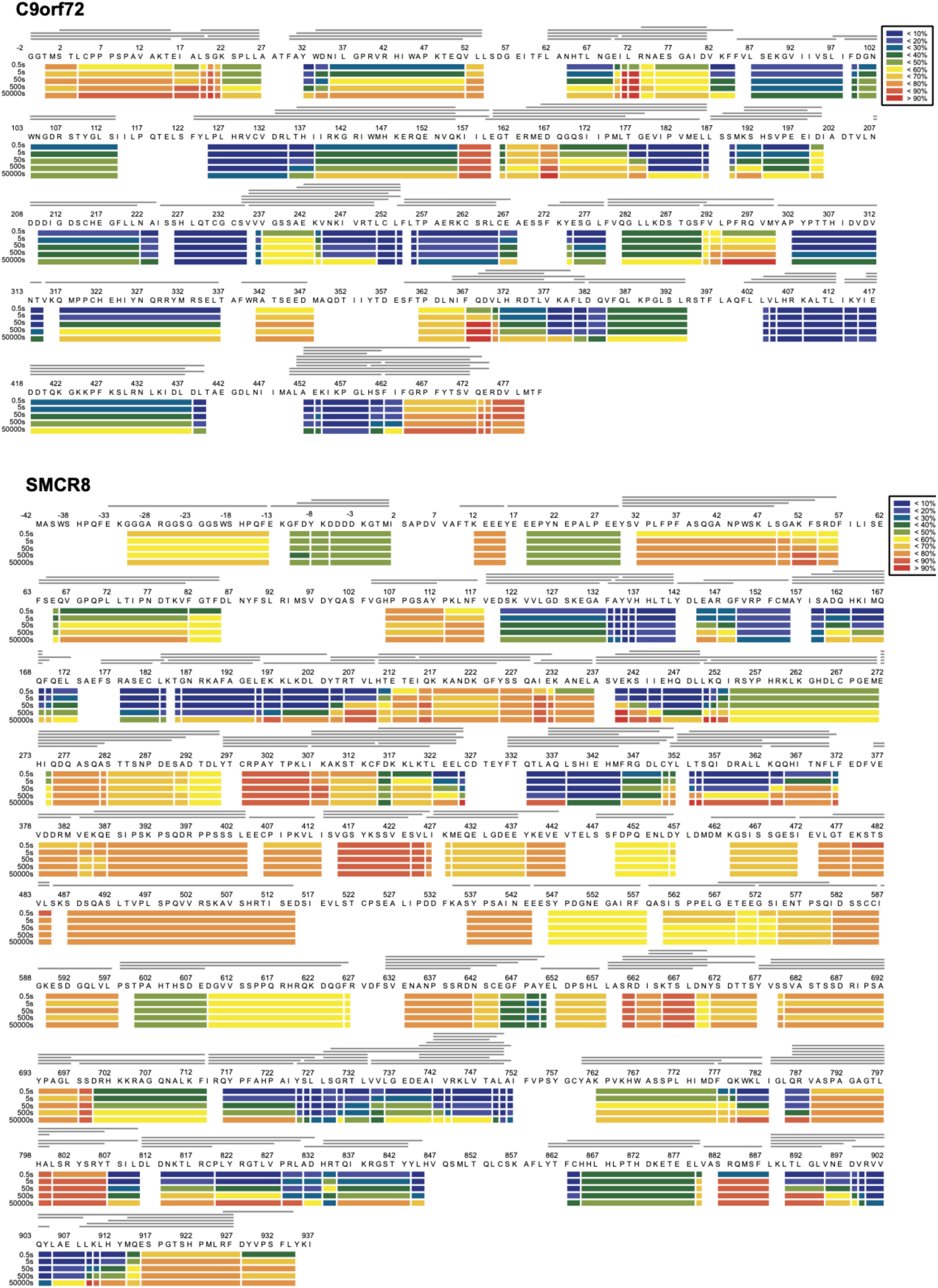
Deuterium uptake of SCARF complex.

**Extended Data Fig. 7:**
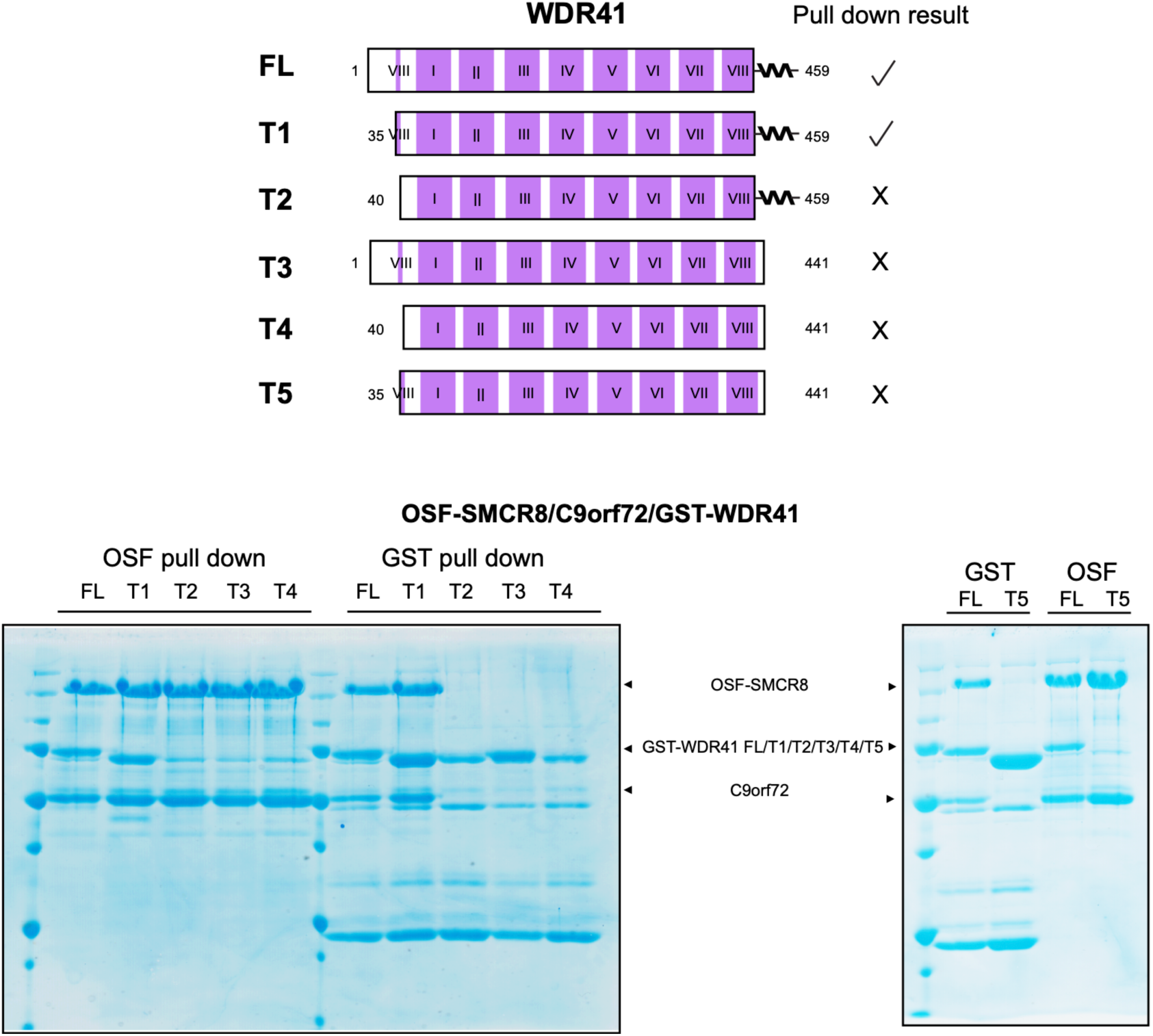
Pull down experiment of WDR41 mutants with SCARF complex.

**Extended Data Fig. 8:**
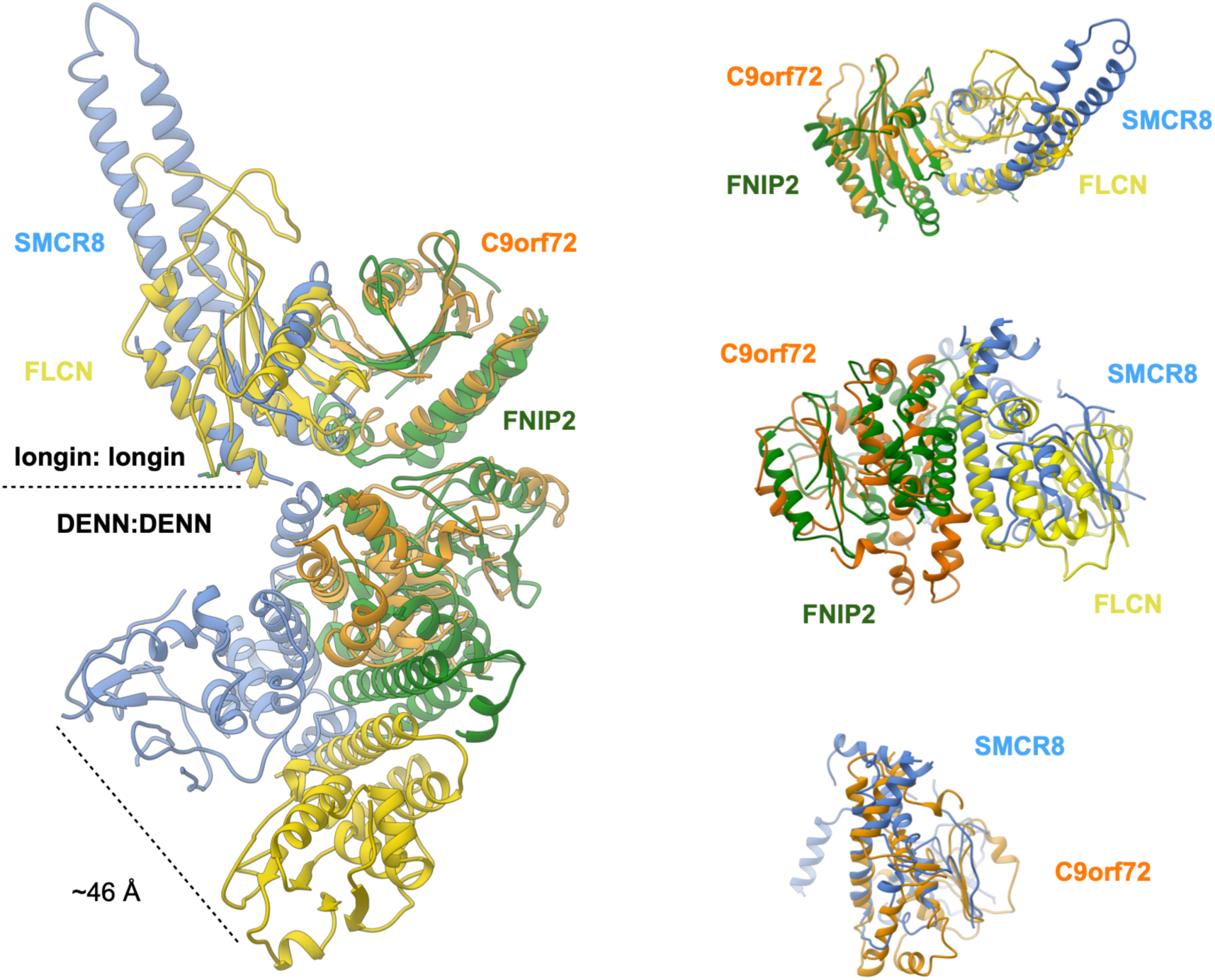
Structural comparison between SCARF and FNIP2-FLCN.

**Extended Data Fig. 9:**
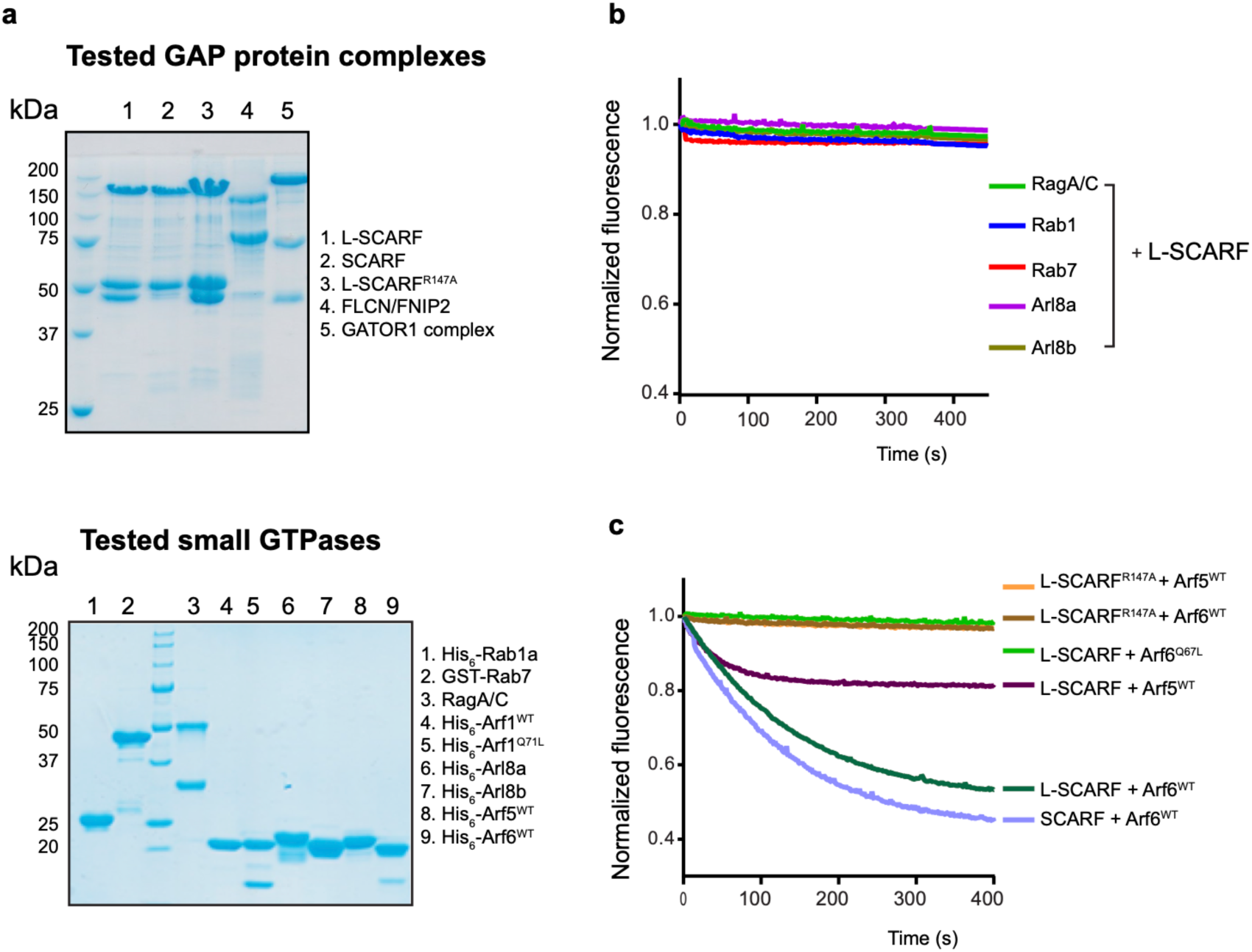
GTPase assay for different small GTPases with L-SCARF complex. **a**, SDS-PAGE of GAP protein complex (top) and GTPase proteins (bottom) used in the experiments. **b**, Tryptophan fluorescence GTPase signal was measured for purified Rag, Arl8a, Arl8b, Rab1a and Rab7 before and after addition of L-SCARF. **c**, Tryptophan fluorescence GTPase signal was measured for purified Arf6^WT or Q67L^ or Arf5^WT^ and before and after addition of SCARF^WT or R147A^-WDR41 or SCARF^WT^.

**Extended Data Fig. 10:**
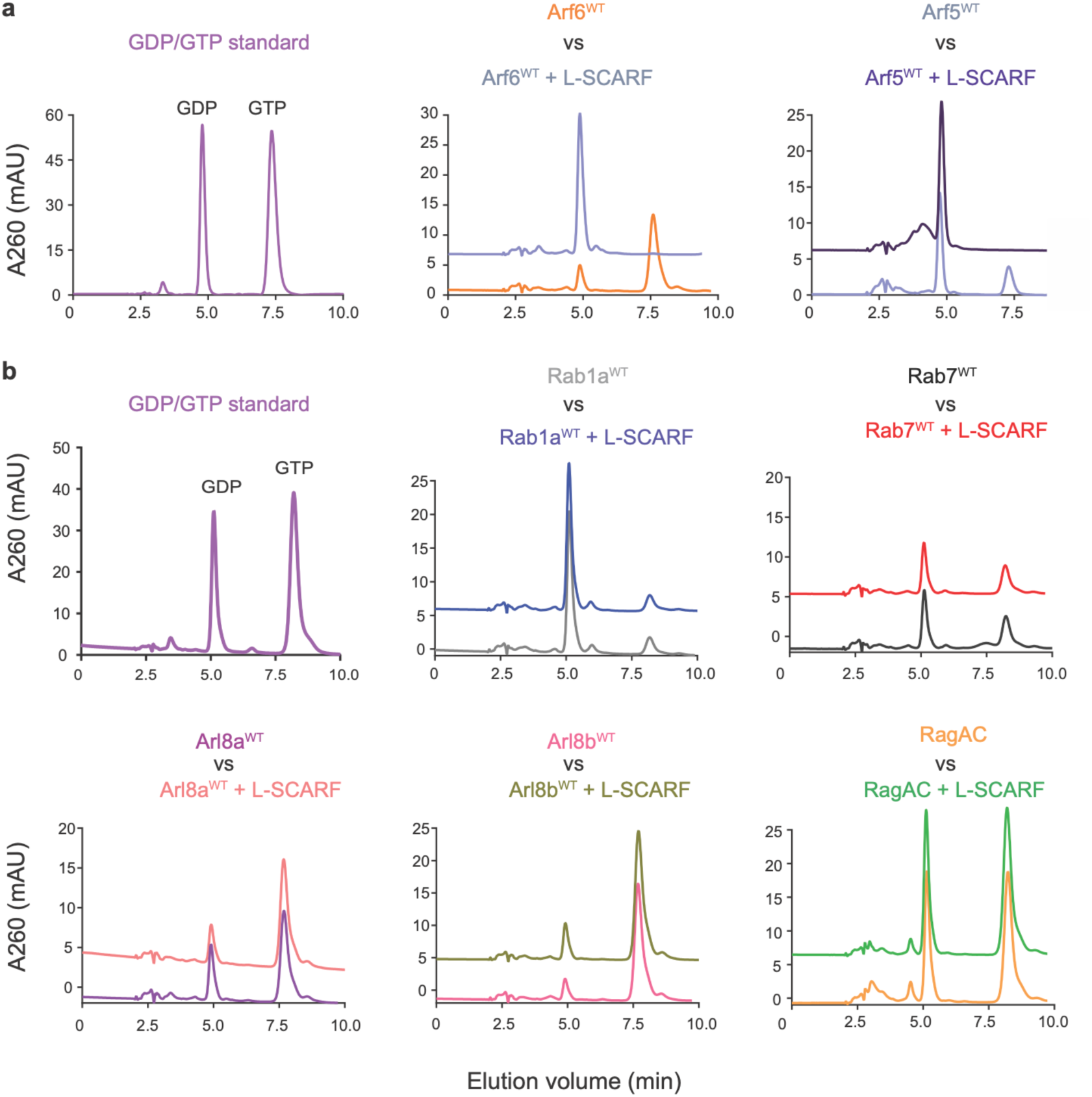
HPLC-based GTPase assay with Arf6, Arf5, Rab1a, Rab7, Arl8a, Arl8b and RagA/C proteins in the absence and addition of L-SCARF complex as indicated.

**Extended Data Table 1.** Cryo-EM data collection, refinement and validation statistics

|  |  |
| --- | --- |
|  | L-SCARF<br>(EMDB-21048)<br>(PDB 6V4U) |
| **Data collection and processing** |  |
| Microscope | Titan Krios |
| Magnification (calibrated) | 43,516 |
| Camera | Quantum-K2 Summit |
| Voltage (kV) | 300 |
| Electron exposure (e<sup>-</sup>/Å<sup>2</sup>) | 59.6 |
| Volta phase plate | Yes |
| Pixel size (Å) | 1.149 |
| Symmetry imposed | C1 |
| Initial particle images (no.) | 4,810,184 |
| Final particle images (no.) | 381,450 |
| Map resolution (Å) | 3.80 |
| FSC threshold | 0.143 |
| **Refinement** |  |
| Initial model used (PDB code) | - |
| Map sharpening *B* factor (Å<sup>2</sup>) | -50 |
| Model composition |  |
| Non-hydrogen atoms | 6817 |
| Protein residues | 1119 |
| Ligands | 0 |
| R.m.s. deviations |  |
| Bond lengths (Å) | 0.002 |
| Bond angles (°) | 0.443 |
| Validation |  |
| MolProbity score | 1.75 |
| Clashscore | 6.07 |
| Poor rotamers (%) | 0 |
| Ramachandran plot |  |
| Favored (%) | 93.79 |
| Allowed (%) | 6.21 |
| Disallowed (%) | 0.00 |

**Extended Data Dataset S1.** HDX Data Summary (L-SCARF and SCARF)

| **Extended Data Dataset S1. HDX Data Summary (L-SCARF and SCARF)** |  |  |
| --- | --- | --- |
| Data Set | L-SCARF | SCARF |
| # of Peptides | SMCR8: 178 | SMCR8: 178 |
|  | C9orf72: 90 | C9orf72: 90 |
|  | WDR41: 73 |  |
| Sequence coverage (%) | SMCR8: 88.8 | SMCR8: 88.8 |
|  | C9orf72: 87 | C9orf72: 87 |
|  | WDR41: 80.4 |  |
| Redundancy | SMCR8: 2.8 | SMCR8: 2.9 |
|  | C9orf72: 3.1 | C9orf72: 3.2 |
|  | WDR41: 2.1 |  |
| Average standard deviation (%) | SMCR8: 0.84 | SMCR8: 0.98 |
|  | C9orf72: 0.88 | C9orf72: 0.91 |
|  | WDR41: 0.85 |  |
| Deuterium time course analyzed (sec) | 0.5/5/50/500/50,000 | 0.5/5/50/500/50,000 |
| Control samples (sec) | 50,000 | 50,000 |
| Replicates | Triplicates/Quadruplicate | Triplicates/Quadruplicate |
| Back-exchange (mean) | SMCR8: 32.23 | SMCR8: 31.03 |
|  | C9orf72: 42.46 | C9orf72: 42.69 |
|  | WDR41: 37.20 |  |

